# Salt Stress Shifts *Isatis indigotica* to Prioritize Defense over Goitrin Biosynthesis

**DOI:** 10.64898/2026.10.13.758776

**Authors:** Lijun Hou, Yuetao Li, Yanyan Zhang, Hua Yao, Yueqiang Zhang

## Abstract

Soil salinization threatens the cultivation quality of *Isatis indigotica*, a medicinal plant whose root bioactivity depends on the glucosinolate-derived compound (R,S)-goitrin. Although moderate abiotic stress is widely assumed to stimulate secondary metabolite accumulation, the temporal dynamics of goitrin biosynthesis under salt stress remain poorly understood. This study quantified goitrin content across four NaCl concentrations (100 – 400 mmol/L) and four time points (2, 6, 12, and 24 h). Time-series transcriptome sequencing was performed under 100 mmol/L NaCl. Contrary to prevailing assumptions, goitrin content declined progressively from 2 to 12 h before partially recovering at 24 h, indicating that salt stress transiently suppresses goitrin accumulation rather than promoting it. Transcriptomic analysis identified 7 glucosinolate biosynthesis genes whose expression positively correlated with goitrin content (p < 0.05), and 6 salt stress-responsive genes negatively correlated with goitrin content (p < 0.05). These opposing patterns suggest that stress defense is transiently prioritized over biosynthesis, with partial metabolic recovery as plants adapt. These findings reveal a defense-biosynthesis trade-off mechanism in *I. indigotica* roots and provide a transcriptomic framework for understanding how abiotic stress reshapes secondary metabolism in medicinal plants.

**Highlights:** Salt stress redirects metabolic resources from goitrin biosynthesis to defense in Isatis indigotica roots, revealing a transcription-driven defense–biosynthesis trade-off.

**Graphic Abstract:** 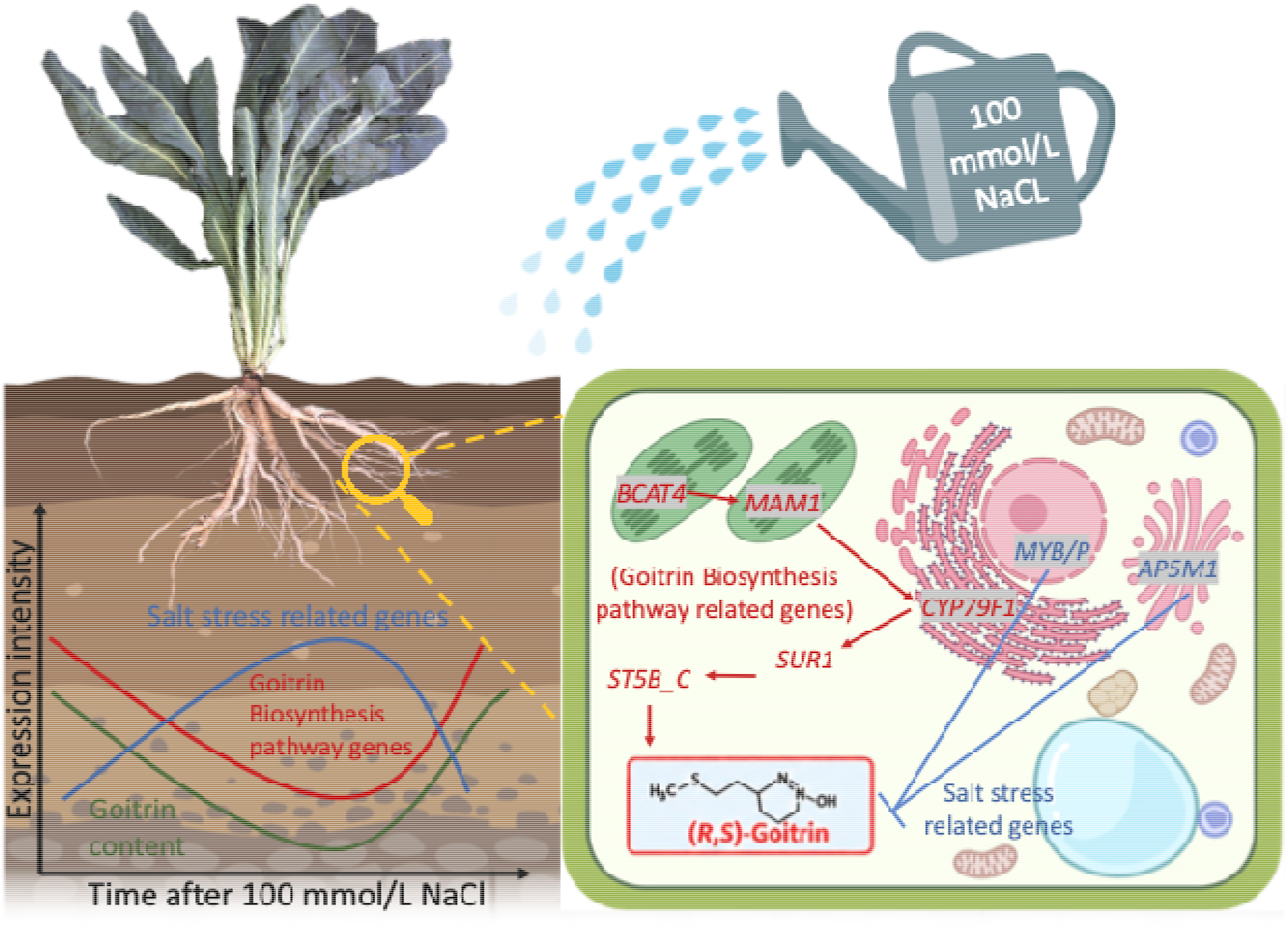

## 1. Introduction

*Isatis indigotica* Fort., a biennial herbaceous plant of the family Brassicaceae, has been widely used in traditional Chinese medicine for over two millennia. Its dried root (Banlangen) exhibits diverse pharmacological activities, including antiviral, antibacterial, anti-inflammatory, and immunomodulatory effects, which have been supported by recent pharmacological and mechanistic studies (Min et al., 2022; Yang et al., 2022).

Soil salinization represents a major global threat to agricultural productivity, affecting more than 20% of irrigated land and posing risks to over one billion hectares worldwide (FAO, 2021). In China, salinized soils are particularly prevalent in the arid and semi-arid regions of the northwest, including Xinjiang, where over 30% of farmland is affected by varying degrees of salinity (Hou et al., 2022). As *I. indigotica* cultivation is increasingly concentrated in these regions to meet growing market demand, soil salinity has emerged as a critical factor influencing both plant growth and root quality. Salt stress imposes ionic and osmotic constraints on plants, disrupting water uptake, ion homeostasis, and normal metabolic processes (Xu et al., 2021). Notably, moderate salt stress has been reported to stimulate secondary metabolite accumulation in several medicinal plants; for example, glycyrrhizin levels increase in *Glycyrrhiza uralensis* under salinity, and alkaloid and flavonoid contents are elevated in *I. indigotica* following NaCl treatment (Bi et al., 2022; Jiang et al., 2022). These findings raised the question of whether similar stimulatory effects on secondary metabolite accumulation might occur in *I. indigotica* roots under salt stress. Among the bioactive metabolites of *I. indigotica* roots, (R,S)-goitrin is the most representative quality marker, officially listed in the Chinese Pharmacopoeia as the standard for root quality assessment (Chinese Pharmacopoeia Commission, 2020). Goitrin is also a glucosinolate hydrolysis product derived from progoitrin and epiprogoitrin via endogenous myrosinase activity, with documented antiviral efficacy against influenza viruses and a role in modulating pro-inflammatory cytokine production (Chen et al., 2021). Whether salt stress modulates goitrin accumulation in *I. indigotica* roots specifically, and the underlying transcriptional mechanisms governing this response, remain largely uncharacterized.

Glucosinolates are a class of sulfur-containing secondary metabolites characteristic of the Brassicaceae family, and their biosynthesis has been most extensively characterized in the model plant *Arabidopsis thaliana* (Chroston et al., 2024). The biosynthetic pathway proceeds through three main stages: amino acid chain elongation, core glucosinolate structure formation, and secondary side-chain modification. In the aliphatic glucosinolate branch, *BCAT4* catalyzes the initial transamination step of methionine chain elongation, *MAM1* mediates condensation reactions during side-chain elongation, cytochrome P450 monooxygenase *CYP79F1* converts elongated methionine derivatives into the corresponding aldoximes, *SUR1* catalyzes S-alkyl-thiohydroximates into thiohydroximates, *ST5b/ST5c* then sulfates the resulting desulfoglucosinolates in the final step of core structure biosynthesis (Choi et al., 2026; Qin et al., 2023; Zhang and Zhang, 2022). While these genes have been functionally characterized in *Arabidopsis*, their roles in *I. indigotica* and their transcriptional responses to salt stress have not been investigated.

Transcriptome sequencing combined with time-series expression profiling has become an effective strategy for identifying candidate genes involved in secondary metabolite biosynthesis in medicinal plants under stress conditions (Wang et al., 2025). Integration of KEGG pathway-based candidate gene screening with Pearson correlation analysis between gene expression dynamics and metabolite accumulation has been applied to identify key biosynthetic genes in medicinal plants under stress conditions (Wang et al., 2024). Transcription factors, including *MYB* play key regulatory roles in glucosinolate biosynthesis and stress-responsive secondary metabolism in Brassicaceae species (Mitreiter and Gigolashvili, 2021). *R2R3-MYB* transcription factors and *MYB*-related proteins are well-established regulators of plant salt stress responses, with multiple family members shown to be rapidly and strongly induced by NaCl treatment in Brassicaceae species (Li et al., 2022; Wang et al., 2021). Nevertheless, whether salt stress responses in *I. indigotica* roots come at the expense of goitrin accumulation remains entirely unknown.

In this study, we investigated the effects of NaCl stress on goitrin accumulation in *I. indigotica* roots across multiple concentrations and time points and performed time-series transcriptome analysis to identify structural genes and transcription factors associated with salt stress-responsive goitrin biosynthesis. We hypothesized that salt stress would transiently suppress goitrin biosynthesis rather than promote it, as the plant redirects metabolic resources toward stress defense during the acute response phase, and that time-series transcriptome analysis would reveal the underlying transcriptional basis of this dynamic. Our findings provide the first transcriptomic evidence linking salt stress signaling to glucosinolate pathway regulation in *I. indigotica* and offer molecular targets for quality-oriented cultivation strategies in saline environments.

## 2. Materials and Methods

### 2.1 Plant material and growth conditions

Seeds of *Isatis indigotica* Fort. from Inner Mongolia origin were used in this study. Seeds were surface-sterilized with 75% ethanol for 30 s and 1% NaClO for 10 min, rinsed thoroughly with sterile distilled water, and germinated on moist filter paper at 25°C in the dark. Seedlings were subsequently transferred to a controlled growth chamber under a 16 h light/8 h dark photoperiod at 25°C/20°C (day/night) with a relative humidity of 60–70% and cultivated in Hoagland nutrient solution until the two-month-old seedling stage before salt stress treatment.

### 2.2 Salt stress treatments

To determine the optimal NaCl concentration for goitrin accumulation, two-month-old seedlings were treated with NaCl solutions at four concentrations (100, 200, 300, and 400 mmol/L) for four durations (2, 6, 12, and 24 h). Control plants were maintained in distilled water under identical conditions. At each time point, roots were harvested, and immediately frozen in liquid nitrogen for subsequent analysis. Each treatment consisted of three independent biological replicates. Based on goitrin quantification results and plant growth assessment, 100 mmol/L NaCl was selected for transcriptome sequencing, as it best maintained plant viability while eliciting a measurable stress response.

### 2.3 Sample preparation and quantification of (R,S)-goitrin by UPLC-MS/MS

The content of (R,S)-goitrin in *I. indigotica* roots was determined by ultra-performance liquid chromatography-tandem mass spectrometry (UPLC-MS/MS)( Agilent Technologies, Santa Clara, CA, USA). Root samples were ground to a fine powder in liquid nitrogen, and 1 g of powder was transferred into a 10 mL centrifuge tube. Eight milliliters of 80% methanol was added, followed by vortexing for 1 min, grinding at 65 Hz for 5 min, and ultrasonication at 4°C for 90 min. The mixture was then allowed to stand at 4°C for 60 min before centrifugation at 12,000 × g for 10 min at 4°C. The supernatant was collected into a new 2 mL centrifuge tube, and 1.5 mL of the extract was passed through a 0.22 μm membrane filter using a disposable sterile syringe. The filtrate was transferred into a headspace vial for UPLC-MS/MS analysis.

Chromatographic separation was performed on an Agilent Eclipse Plus C18 column (1.8 μm, 2.1 mm × 100 mm) with a column temperature of 40°C, a flow rate of 0.300 mL/min, and an injection volume of 2 μL. The mobile phase consisted of 0.1% formic acid in water (A) and acetonitrile (B), with gradient elution as described in **Table S1**. Mass spectrometry detection was performed using an electrospray ionization (ESI) source operated at 300°C with a gas flow rate of 7 L/min, nebulizer pressure of 15 psi, capillary voltage of 4 kV, sheath gas temperature of 250°C, sheath gas flow rate of 11 L/min, and nozzle voltage of 1.5 kV. Analytes were detected in multiple reaction monitoring (MRM) mode. A calibration curve was constructed by plotting peak area against analyte concentration using a series of mixed standard solutions, with the calibration curve showing a coefficient of determination (R² > 0.99) as described in **Method S1 and Table S2**. (R,S)-goitrin content was expressed as mg/g dry weight, and all measurements were performed in triplicate.

### 2.4 RNA extraction and transcriptome sequencing

Total RNA was extracted from frozen *I. indigotica* root samples at Shanghai Majorbio Bio-pharm Technology Co., Ltd. (Shanghai, China) using TRIzol® Reagent. RNA quality was assessed using a 5300 Bioanalyzer (Agilent Technologies, Santa Clara, CA, USA) and NanoDrop ND-2000 spectrophotometer, and only high-quality samples (OD□□□/□□□ = 1.8–2.2, RQN ≥ 6.5 (equivalent to RIN), ≥ 1 μg) were used for library construction. A total of 24 strand-specific RNA-seq libraries were constructed from 100 mmol/L NaCl-treated and control root samples at four time points (2, 6, 12, and 24 h), with three biological replicates per treatment (4 time points × 2 treatments × 3 replicates). Libraries were prepared following the Illumina® Stranded mRNA Prep Ligation protocol and sequenced on the Illumina NovaSeq platform (2 × 150 bp paired-end reads) at Shanghai Majorbio Bio-pharm Technology Co., Ltd. Each sample generated at least 5.91 Gb of clean data with a Q30 base percentage ≥ 95.12%. The raw sequencing data have been deposited in the NCBI Sequence Read Archive (SRA) under BioProject accession number PRJNA1455832.

### 2.5 Transcriptome data processing and differential expression analysis

Raw reads were quality-controlled using fastp and aligned to the *I. indigotica* reference genome (https://www.bic.ac.cn/IMP/#/speciesInfo?ID=255) using HISAT2, with mapping rates ranging from 74.55% to 82.51%. Mapped reads were assembled using StringTie in a reference-based approach. Gene expression levels were quantified using RSEM and expressed as transcripts per million reads (TPM). DEGs between NaCl-treated and control samples at each time point were identified using DESeq2 with criteria of |log□ FC| ≥ 1.322 (corresponding to an approximately 2.5-fold change threshold) and padjust < 0.01. KEGG pathway enrichment analysis was performed using KOBAS (corrected P ≤ 0.05). All bioinformatic analyses were performed using the Majorbio Cloud platform (www.majorbio.com) (Han et al., 2024).

### 2.6 KEGG Pathway-Based Candidate Gene Screening and Correlation Analysis

To identify structural genes potentially associated with goitrin biosynthesis under salt stress, candidate genes were screened based on KEGG pathway annotation. Differentially expressed genes (DEGs) annotated to the glucosinolate biosynthesis pathway (map00966) were extracted from the transcriptome dataset. Pearson correlation analysis was then performed between the mean expression level (E mean, calculated from TPM values across three biological replicates) of each candidate gene and the mean (R,S)-goitrin content at four time points (2, 6, 12, and 24 h) under 100 mmol/L NaCl treatment. Genes showing significant positive correlation with goitrin accumulation (r > 0, p < 0.05) were retained as key candidate structural genes.

To identify salt stress-responsive genes whose expression dynamics were negatively associated with goitrin accumulation, DEGs functionally annotated as salt stress-related were retrieved from the transcriptome dataset based on KEGG annotation. Pearson correlation analysis was performed between the mean expression level of each salt stress-related DEG and goitrin content across the four time points, and genes showing significant negative correlation (r < 0, p < 0.05) were identified. Correlation coefficients (r) and p-values were calculated using the SciPy library in Python (v3.9), and scatter plots with linear regression lines were generated using Matplotlib.

### 2.7 Statistical analysis

Statistical differences in (R,S)-goitrin content between NaCl-treated and control groups at each time point were assessed by Student’s t-test using GraphPad Prism 9, and data are presented as mean ± standard deviation (SD) of three independent biological replicates (n = 3). For Pearson correlation analyses between candidate gene expression and goitrin content, gene expression levels are reported as mean ± standard error of the mean (SEM, n = 3); scatter plots with bilateral SEM error bars and linear regression fits were generated using Python (v3.9; Python Software Foundation, 2021) with the SciPy and Matplotlib libraries. Heatmaps showing log□ FC (NaCl-treated vs. control) across four time points were generated using Python (v3.9; Python Software Foundation, 2021) with the Matplotlib library. Transcriptomic data processing and differential expression analysis were performed using R (v4.1.0; R Core Team, 2021). ChatGPT (OpenAI) was used to assist in generating the pathway diagram in Fig. 6a.

## 3. Results and Discussion

### 3.1 NaCl stress transiently suppresses and partially restores (R,S)-goitrin accumulation in I. indigotica roots

To investigate the effects of salt stress on goitrin accumulation, two-month-old *I. indigotica* seedlings were treated with NaCl at four concentrations (100, 200, 300, and 400 mmol/L) at four time points (2, 6, 12, and 24 h). The results showed that (R,S)-goitrin content was significantly affected by both NaCl concentration and treatment duration (**Fig. 1**).

**Fig. 1.**
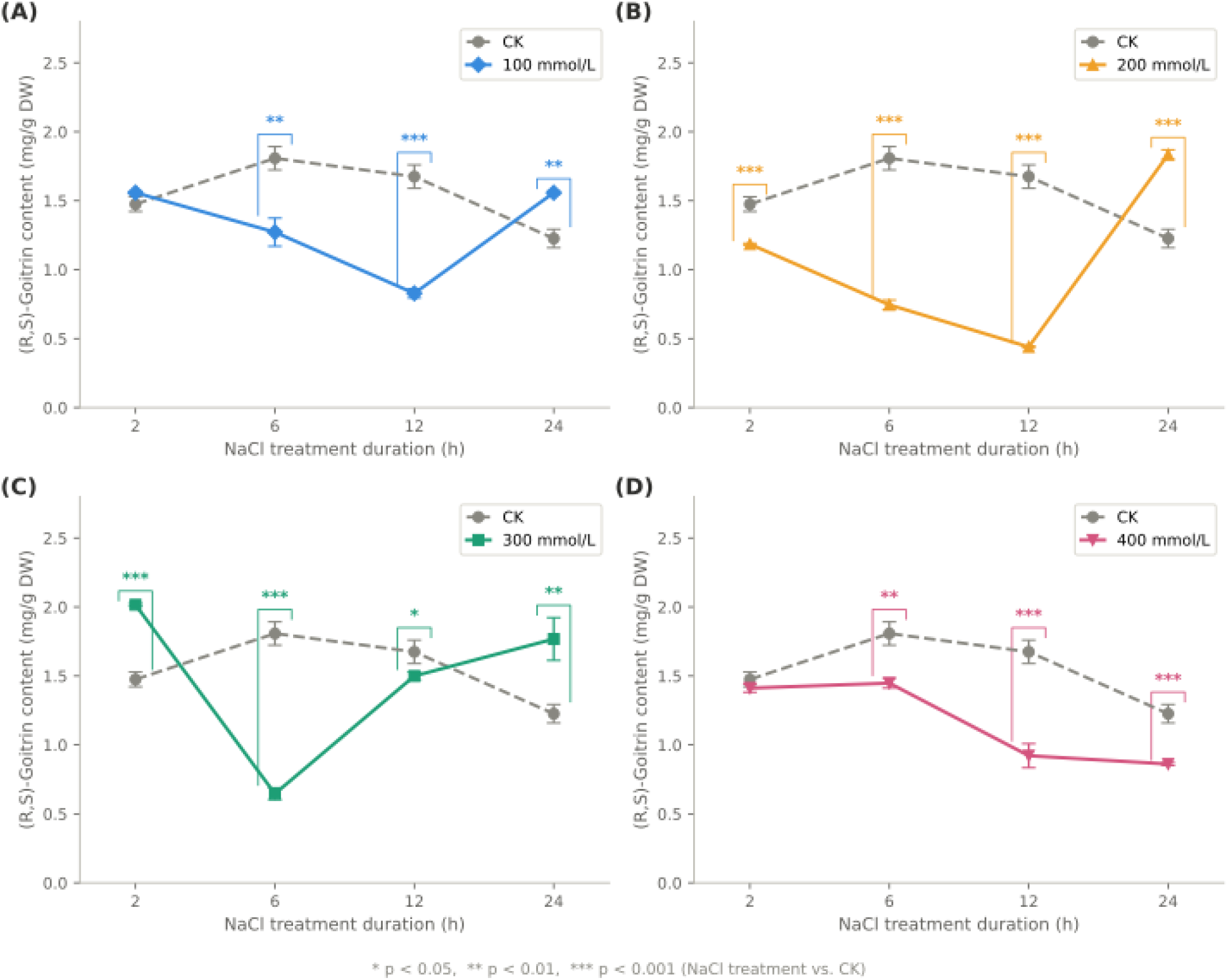
Effects of NaCl stress on (R,S)-goitrin content in *Isatis indigotica* roots. Two-month-old seedlings were treated with A) 100, B) 200, C) 300, or D) 400 mmol/L NaCl for 2, 6, 12, and 24 h. Distilled water-treated plants served as controls (CK). Data are presented as mean ± SD (n = 3). Asterisks indicate significant differences between NaCl-treated and CK groups at the corresponding time point (Student’s t-test: * *p* < 0.05; ** *p* < 0.01; *** *p* < 0.001).

At 100 and 200 mmol/L NaCl, goitrin content declined relative to the corresponding CK from 6 h, reached its lowest point at 12 h, and partially recovered by 24 h. At 100 mmol/L, goitrin content dropped to a minimum of 0.828 ± 0.033 mg/g DW at 12 h before recovering to 1.558 ± 0.010 mg/g DW at 24 h, which was significantly higher than the corresponding CK (1.226 ± 0.066 mg/g DW). At 300 mmol/L NaCl, the suppression was more rapid, with goitrin content falling sharply at 6 h relative to CK before beginning to recover at 12 h and reaching levels above CK by 24 h. At 400 mmol/L NaCl, goitrin content declined progressively across all time points without recovery, suggesting that severe ionic toxicity irreversibly disrupts secondary metabolic capacity rather than causing a temporary reallocation of metabolic resources.

Notably, at 100 mmol/L NaCl, plant growth was not visibly inhibited over 21 days, whereas higher concentrations caused progressive leaf chlorosis, wilting, and eventual necrosis in a concentration-dependent manner (**Fig. 2**). These observations indicate that 100 mmol/L represents a moderate stress level at which plants remain physiologically viable while mounting a measurable stress response, making it the most suitable concentration for subsequent transcriptome analysis.

**Fig. 2.**
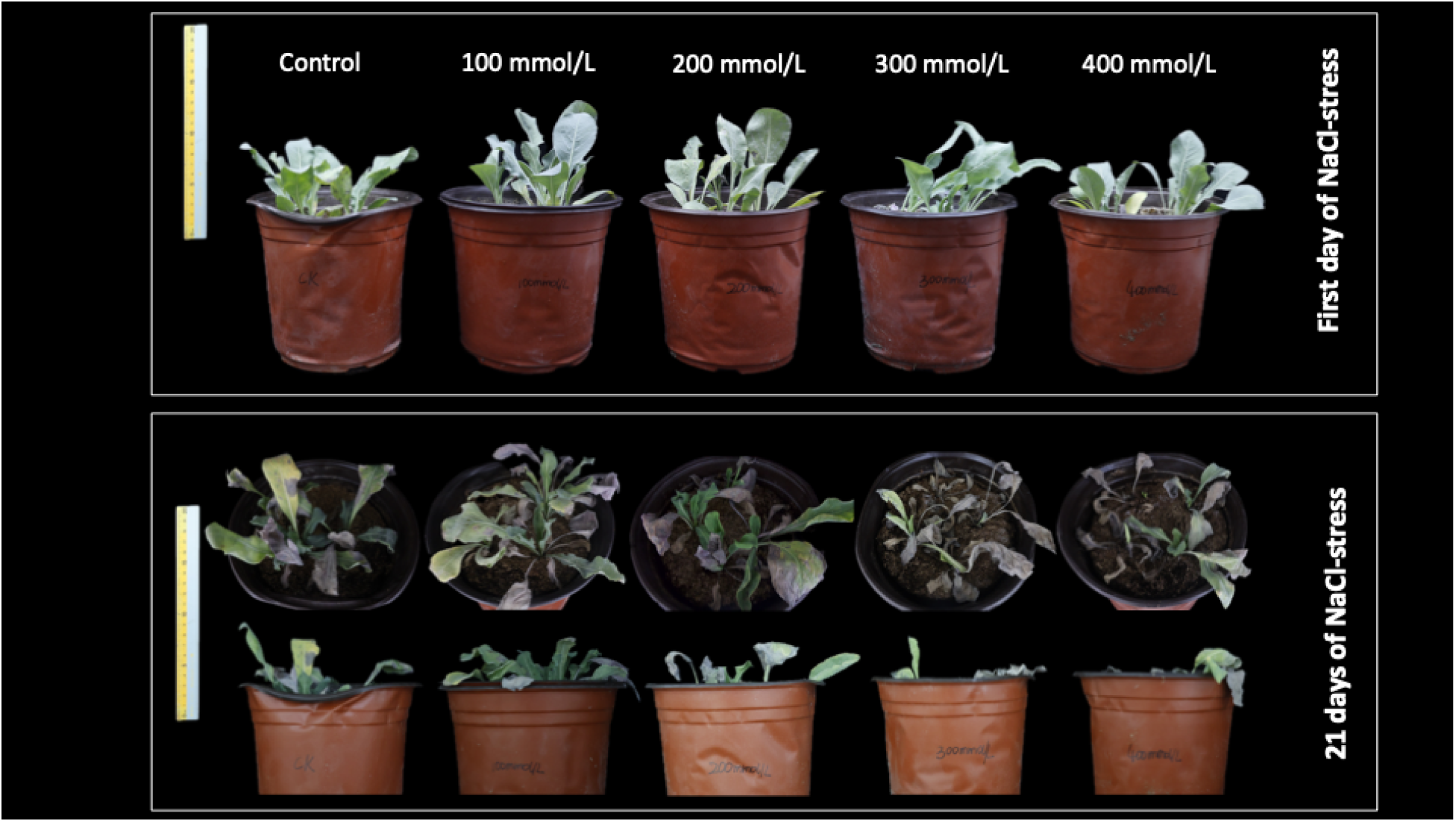
Phenotypic responses of *Isatis indigotica* seedlings to NaCl stress at different concentrations. Two-month-old seedlings were treated with 0 (control), 100, 200, 300, or 400 mmol/L NaCl. Representative photographs were taken on the first day (upper panel) and after 21 days (lower panel) of NaCl treatment. A ruler is shown on the left of each panel for size reference.

Taken together, these results suggest that salt stress does not simply stimulate goitrin accumulation, as has been reported for secondary metabolites in some medicinal plants under moderate abiotic stress (Ullah et al., 2025; Bi et al., 2022; Jiang et al., 2022). Instead, goitrin biosynthesis appears to be transiently suppressed during the early stress response, with partial recovery occurring as plants begin to adapt by 24 h. This pattern is consistent with the hypothesis that plants prioritize defense responses over secondary metabolite biosynthesis under acute salt stress, redirecting metabolic resources toward stress tolerance mechanisms.

### 3.2 Salt stress induces time-dependent transcriptional responses in I. indigotica roots

To identify salt stress-responsive genes in *I. indigotica* roots, RNA-seq was performed on samples collected at four time points (2, 6, 12, and 24 h) under 100 mmol/L NaCl treatment and corresponding controls. A total of 24 libraries generated 159.09 Gb of clean data, with each sample yielding at least 5.91 Gb and a Q30 base percentage ≥ 95.12%, indicating high sequencing quality. Clean reads were aligned to the reference genome with mapping rates ranging from 74.55% to 82.51%.

Differential expression analysis identified a total of 2,919, 9,341, 3,872, and 5,175 DEGs at 2, 6, 12, and 24 h, respectively. At 2 h, 1,367 genes were upregulated and 1,552 were downregulated. The largest number of DEGs was observed at 6 h (4,451 upregulated and 4,890 downregulated), suggesting broad transcriptional reprogramming at this early stress stage. At 12 h, the number of DEGs decreased to 3,872 (1,127 upregulated and 2,745 downregulated), with downregulated genes predominating, consistent with partial stress adaptation. At 24 h, the DEG number rebounded to 5,175, with upregulated genes (3,694) substantially exceeding downregulated genes (1,481), suggesting a shift in transcriptional priorities at this stage. Venn diagram analysis revealed that 205 DEGs were shared across all four time points, representing the core salt-responsive gene set, while the majority of DEGs were time point-specific (**Fig. S1**). The shift from predominantly downregulated genes at 12 h to predominantly upregulated genes at 24 h mirrors the partial recovery of goitrin content at the same time point, suggesting a shift in transcriptional priorities toward biosynthetic activation during the late stress response (**Fig. 3**).

**Fig. 3.**
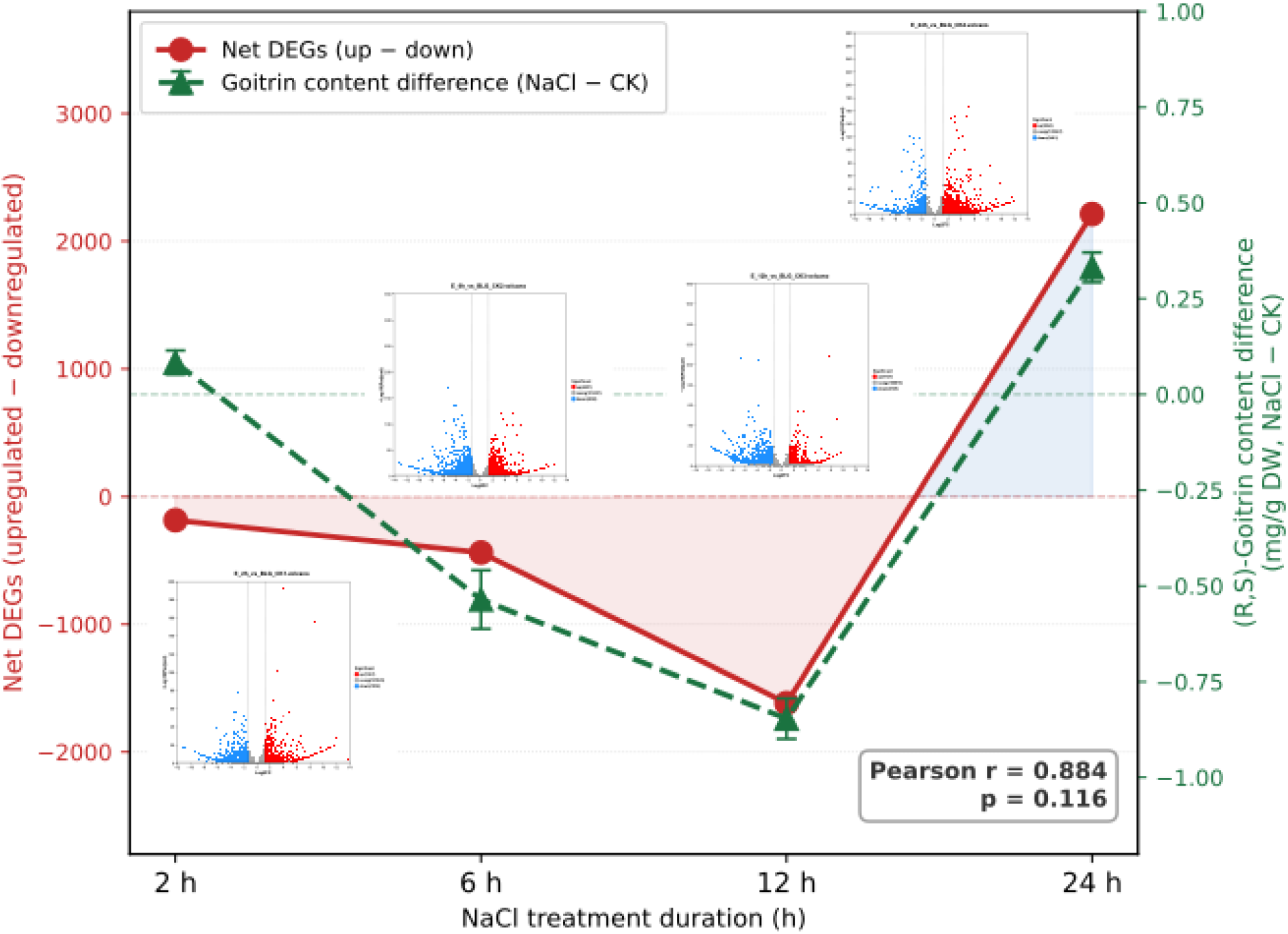
Temporal dynamics of net transcriptional changes and (R,S)-goitrin content in Isatis indigotica roots under 100 mmol/L NaCl stress. The net number of differentially expressed genes (DEGs; upregulated minus downregulated, red line) and the difference in (R,S)-goitrin content between NaCl-treated and control plants (green dashed line) are plotted across four time points (2, 6, 12, and 24 h). Inset volcano plots show DEG distribution at each time point (red: upregulated; blue: downregulated; gray: non-significant). Positive net DEG values indicate predominance of upregulated genes; negative values indicate predominance of downregulated genes. Error bars represent SEM (n = 3). DEGs were identified by DESeq2 (|logß FC| ≥ 1.322, padj < 0.01). Pearson correlation between net DEG balance and goitrin content difference: r = 0.884, p = 0.116.

Similar time-dependent transcriptional responses have been reported in other plants under salt stress, where the number and direction of DEGs shift substantially across treatment durations, reflecting the transition from acute stress signaling to adaptive metabolic responses (Wang et al., 2022; Hu et al., 2022). Notably, the net DEG balance (upregulated minus downregulated genes) showed a temporal trend broadly consistent with the changes in goitrin content relative to control across the four time points, suggesting that the overall direction of transcriptional reprogramming may reflect the dynamic regulation of goitrin biosynthesis under salt stress, though this association did not reach statistical significance given the limited number of time points (n = 4). These findings prompted further investigation into the specific genes and pathways underlying the observed goitrin dynamics.

### 3.3 Glucosinolate biosynthesis genes show expression dynamics consistent with goitrin accumulation under salt stress

#### 3.3.1 Seven glucosinolate biosynthesis genes show significant positive correlation with goitrin accumulation under salt stress

To identify structural genes potentially associated with goitrin accumulation under salt stress, all DEGs annotated to the glucosinolate biosynthesis pathway (KEGG map00966) were extracted from the transcriptome dataset. A total of 42 DEGs were mapped to this pathway under 100 mmol/L NaCl treatment (**Fig. S2**), representing approximately 40% of the 105 genes annotated to this pathway in the KEGG database, indicating that salt stress elicits a broad transcriptional response across the glucosinolate biosynthesis pathway rather than affecting isolated steps. Pearson correlation analysis between the mean expression level of each gene and (R,S)-goitrin content across four time points (2, 6, 12, and 24 h) identified seven genes with significant positive correlation (*r* > 0.957, *p* < 0.05; **Fig. 4a**), including two copies of *BCAT4*, two copies of *SUR1*, and one each of *CYP79F1*, *MAM1*, and *ST5B_C*. These seven genes span three sequential stages of the aliphatic glucosinolate biosynthesis pathway: methionine chain elongation (*BCAT4*, *MAM1*), core glucosinolate structure formation (*CYP79F1*, *SUR1*), and sulfation (*ST5B_C*), suggesting that salt stress coordinately affects multiple steps of the pathway in association with goitrin accumulation.

**Fig. 4.**
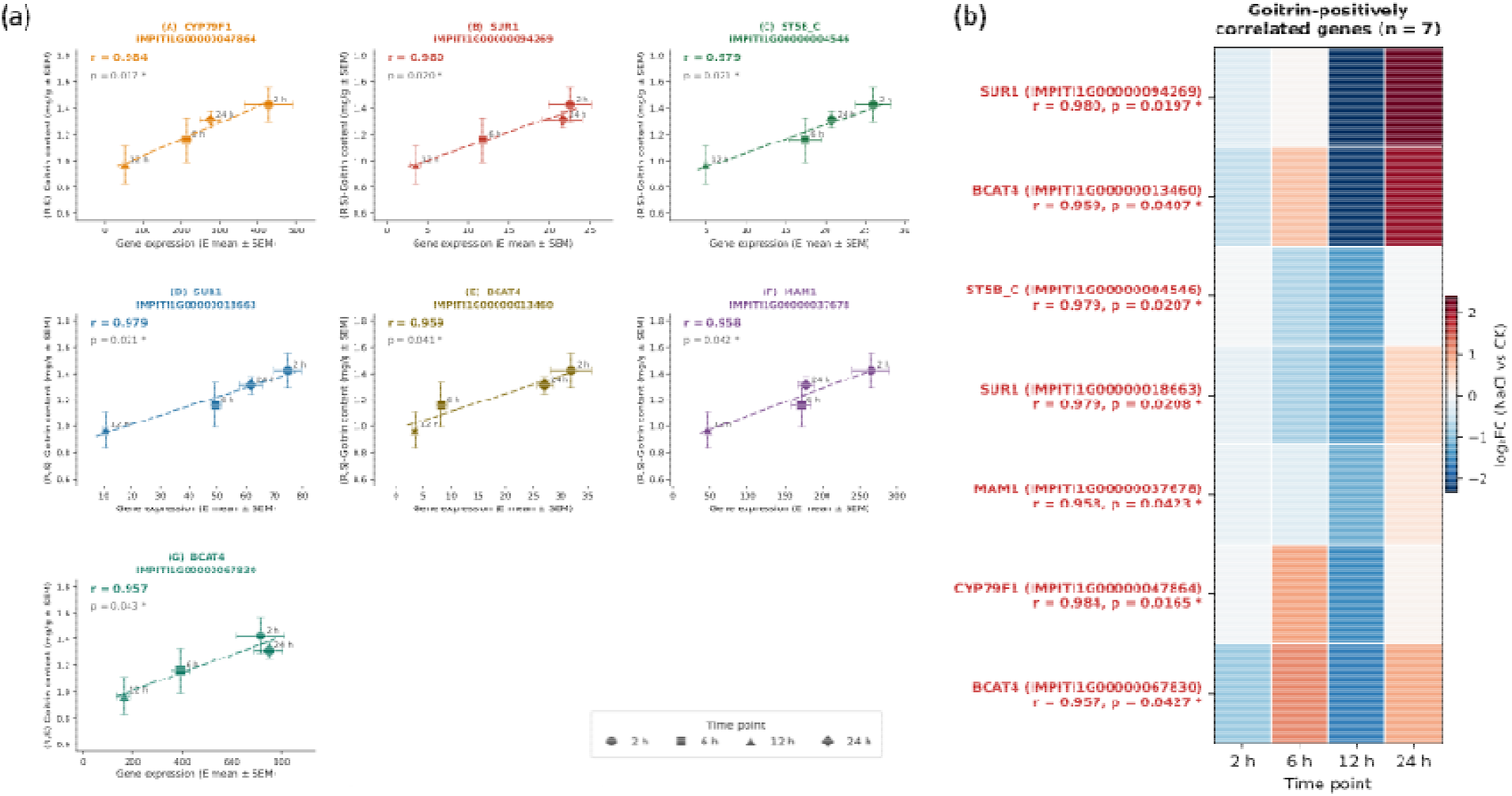
Expression dynamics of seven glucosinolate biosynthesis genes positively correlated with (R,S)-goitrin accumulation in *Isatis indigotica* roots under 100 mmol/L NaCl. **(a)** Scatter plots showing Pearson correlation between gene expression (x-axis) and goitrin content (y-axis) at 2, 6, 12, and 24 h for CYP79F1, SUR1 (×2), ST5B_C, BCAT4 (×2), and MAM1. Dashed lines indicate linear regression fits; r and p-values are shown in each panel. Error bars represent SEM (n = 3). Horizontal and vertical error bars represent SEM of gene expression and goitrin content, respectively (n = 3). * *p* < 0.05; ** *p* < 0.01. **(b)** Heatmap of logß FC (NaCl vs. control) for the seven genes across four time points, ordered by hierarchical clustering. Red: upregulation; blue: downregulation.

#### 3.3.2 All seven candidate genes reach minimum expression at 12 h, mirroring the temporal decline of goitrin content

These temporal correlations were further examined by visualizing log□ FC-based expression dynamics of the seven candidate genes across the four time points (**Fig. 4b**). All seven genes reached their lowest expression levels at 12 h relative to the corresponding controls, coinciding precisely with the lowest goitrin content recorded under 100 mmol/L NaCl treatment (section 3.1). By 24 h, expression of all seven genes partially recovered, paralleling the partial restoration of goitrin content. This coordinated temporal pattern across multiple biosynthetic steps indicates that salt stress does not selectively suppress individual enzymatic reactions, but rather downregulates the glucosinolate biosynthesis pathway as a whole during the acute stress phase, with transcriptional recovery accompanying plant adaptation.

This pathway-wide suppression was further supported by the expression dynamics of the remaining 35 DEGs mapped to map00966 (**Fig. S3**). The majority (19 of 35) followed a pattern broadly similar to the seven positively correlated genes, showing downregulation at 12 h followed by partial or full recovery at 24 h, further corroborating the notion of pathway-wide suppression during peak salt stress with subsequent transcriptional recovery. Nine genes showed sustained downregulation throughout the 24 h observation window, suggesting that certain biosynthetic steps may require longer recovery times or remain persistently suppressed under continuous salt stress. The remaining seven genes displayed mixed or oscillatory expression patterns, potentially reflecting more complex regulatory dynamics or cross-pathway interactions that warrant further investigation. Collectively, the expression dynamics of all 42 DEGs mapped to map00966 reinforce the conclusion that salt stress broadly perturbs glucosinolate biosynthesis at the transcriptional level, with the magnitude and reversibility of suppression varying across individual genes.

#### 3.3.3 The seven candidate genes span all three biosynthetic stages and show functional redundancy at key steps

The functional roles of the seven positively correlated genes are consistent with their established positions in the aliphatic glucosinolate biosynthesis pathway. In the chain elongation stage, *BCAT4* catalyzes the initial transamination of methionine, and *MAM1* mediates the iterative condensation reactions that determine glucosinolate side-chain length and yield in Brassicaceae (Kitainda et al., 2024). The identification of two gene copies each for *BCAT4* and *SUR1* with nearly identical correlation coefficients suggests functional redundancy at these two biosynthetic steps, a phenomenon previously reported in *Arabidopsis thaliana* where multiple *BCAT* paralogs contribute to methionine chain elongation (Zhao et al., 2026). In the core structure formation stage, *CYP79F1* encodes a cytochrome P450 monooxygenase that converts elongated methionine derivatives into the corresponding aldoximes, a committed and rate-limiting step in aliphatic glucosinolate biosynthesis (Jhingan et al., 2023), while *SUR1* catalyzes the subsequent C–S lyase reaction to form thiohydroximate. Finally, in the sulfation stage, *ST5B_C* encodes a sulfotransferase responsible for converting desulfoglucosinolates into mature glucosinolates (Qin et al., 2023). The coordinated downregulation of genes spanning all three stages further supports the notion of pathway-wide suppression rather than selective inhibition of individual enzymatic steps under salt stress. Whether this transcriptional suppression of the glucosinolate biosynthesis pathway is coupled with the concurrent activation of salt stress defense genes was examined in section 3.4.

### 3.4 Salt stress-responsive genes show expression dynamics inversely associated with goitrin accumulation

#### 3.4.1 Six salt stress-responsive genes show significant negative correlation with goitrin accumulation under salt stress

To investigate whether salt stress defense responses are temporally associated with goitrin suppression, DEGs functionally annotated to salt stress response via KEGG were subjected to Pearson correlation analysis against (R,S)-goitrin content across four time points (2, 6, 12, and 24 h). Six genes showed significant negative correlation with goitrin content (*r* < −0.969, *p* < 0.05; **Fig. 5a**), comprising three *MYB* transcription factors, two *MYB*-related proteins (*MYBP*), and one adaptor protein complex subunit (*AP5M1*). The predominance of *MYB* and *MYB*-related proteins among these negatively correlated genes suggests that transcription factor-mediated salt stress signaling may play a role in suppressing goitrin biosynthesis during the early stress response.

**Fig. 5.**
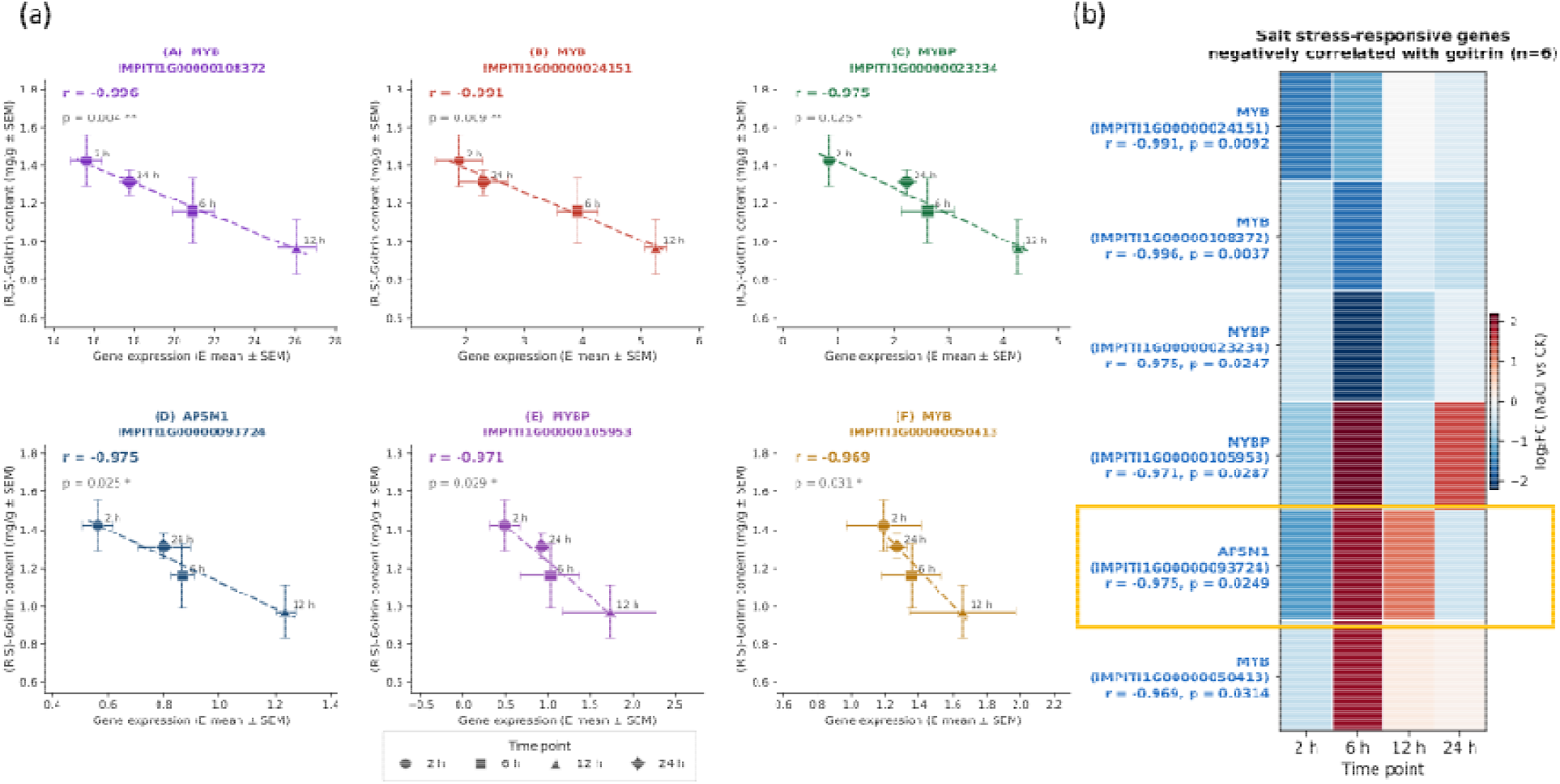
Expression dynamics of six salt stress-responsive genes negatively correlated with (R,S)-goitrin accumulation in *Isatis indigotica* roots under 100 mmol/L NaCl treatment. **(a)** Scatter plots showing Pearson correlation between gene expression (x-axis) and goitrin content (y-axis) at 2, 6, 12, and 24 h for MYB (×3), MYBP (×2), and AP5M1. Dashed lines indicate linear regression fits; r and p-values are shown in each panel. Error bars represent SEM (n = 3). Horizontal and vertical error bars represent SEM of gene expression and goitrin content, respectively (n = 3). * *p* < 0.05; ** *p* < 0.01. **(b)** Heatmap of logß FC (NaCl vs. control) for the six genes across four time points, ordered by hierarchical clustering. The orange box highlights AP5M1, which showed peak upregulation at 6 h coinciding with the lowest goitrin content. Red: upregulation; blue: downregulation.

#### 3.4.2 Salt stress-responsive genes peak at early time points and decline as goitrin content begins to recover

Log□ FC-based expression dynamics of the six candidate genes were visualized across the four time points (**Fig. 5b**). Among these, three genes (*MYBP* IMPITI1G00000105953, *MYB* IMPITI1G00000050413, and *AP5M1* IMPITI1G00000093724) showed marked upregulation at 6 h (log□FC > 1.9), coinciding with the period of greatest goitrin suppression, and subsequently declined toward baseline by 24 h as goitrin content began to recover. This pattern is directly consistent with a stress defense activation model, in which early induction of defense-related genes corresponds temporally with suppression of secondary metabolite biosynthesis. *MYB* (IMPITI1G00000108372) and *MYB* (IMPITI1G00000024151) also exhibited upward trends at 12 h relative to earlier time points, further supporting stress-responsive induction during the acute phase. The remaining gene, *MYBP* (IMPITI1G00000023234), showed downregulation relative to control throughout the observation window. However, its absolute expression level still peaked at 12 h, coinciding with the lowest goitrin content, thus maintaining a significant negative correlation with goitrin accumulation. Although this pattern does not conform to classical stress-induction dynamics, it may reflect the capacity of certain *MYB* members to act as transcriptional repressors of secondary metabolite biosynthesis under salt stress (Dang et al., 2026), or alternatively, the complex and heterogeneous transcriptional reprogramming that characterizes the early salt stress response, in which both up- and downregulation of stress-associated transcription factors have been widely observed (Gan et al., 2024; Basharat et al., 2026).

Despite this divergence in directionality, all six genes shared a common feature that their expression dynamics were inversely correlated with goitrin content across the four time points, with the lowest goitrin levels at 12 h corresponding to the most divergent expression relative to the biosynthesis genes, and partial recovery of goitrin at 24 h accompanying a return of all six genes toward control levels. These opposing temporal dynamics thus provide partial but convergent transcriptomic evidence linking salt stress-responsive gene activity to the suppression of goitrin accumulation.

#### 3.4.3 MYB-dominated stress response genes are functionally consistent with salt defense roles, while AP5M1 represents a novel candidate

The predominance of *R2R3-MYB* transcription factors and *MYB*-related proteins among the negatively correlated genes is consistent with their well-established roles as regulators of plant salt stress responses (Bhatt et al., 2025; Hölzl et al., 2023). *R2R3-MYB* family members have been shown to be rapidly and strongly induced by NaCl treatment across diverse plant species, where they coordinate ion homeostasis, reactive oxygen species scavenging, and osmotic adjustment (Zhang et al., 2026; Ma et al., 2025). The strong negative correlation between *MYB/MYBP* expression and goitrin content observed here suggests that the early activation of these stress defense regulators may compete with or suppress the transcriptional activity of glucosinolate biosynthesis genes, consistent with the broader concept of resource reallocation from secondary metabolism to primary stress defense under acute abiotic stress.

The functional role of *AP5M1* in this context remains unclear, as adaptor protein complex subunits have not been directly implicated in secondary metabolite regulation. Notably, however, *AP5M1* exhibited the most temporally precise inverse pattern among the six negatively correlated genes. It was downregulated at 2 h (log□FC = −1.26), reached peak upregulation at 6 h (log□FC = +1.98), coinciding precisely with the lowest goitrin content recorded across the observation window, and subsequently declined progressively at 12 h (log□FC = +1.20) and 24 h (log□FC = −0.47), paralleling the partial recovery of goitrin accumulation. This temporally precise inverse relationship suggests that whatever its mechanistic role, *AP5M1* activity is tightly coupled to the suppression and recovery of goitrin biosynthesis under salt stress.

*AP5M1* encodes the μ subunit of adaptor protein complex 5 (AP-5), which mediates intracellular cargo sorting and vesicle trafficking between late endosomes and lysosomes (Hirst et al., 2018). While AP-5-mediated endomembrane trafficking has been implicated in cellular homeostasis during salinity stress in plants (Baena et al., 2024), the function of *AP5M1* in plant secondary metabolite regulation remains entirely uncharacterized, the predicted partial conservation of AP-5 in plant genomes has yet to be functionally validated (Wang et al., 2023). Its negative correlation with goitrin content may reflect a broader restructuring of intracellular trafficking under salt stress conditions, and to our knowledge, this is the first report implicating an AP-5 subunit in the context of salt stress-responsive secondary metabolism in a medicinal plant, representing a potentially novel regulatory node that warrants further functional investigation.

### 3.5 Salt stress transiently suppresses goitrin biosynthesis by prioritizing defense over secondary metabolism

The integrated analysis of glucosinolate biosynthesis genes and salt stress-responsive genes reveals a coherent mechanistic model of how salt stress regulates goitrin accumulation in *I. indigotica* roots. The seven positively correlated glucosinolate biosynthesis genes identified in this study span all three sequential stages of the aliphatic glucosinolate biosynthesis pathway: *BCAT4* and *MAM1* in the chain elongation stage (chloroplast), *CYP79F1* and *SUR1* in the core structure formation stage (endoplasmic reticulum and cytoplasm), and *ST5B_C* in the sulfation stage (cytoplasm), covering the complete biosynthetic route from methionine to progoitrin and ultimately to (R,S)-goitrin (Zhu et al., 2024; Zhang et al., 2022; **Fig. 6a**). This pathway-wide distribution indicates that salt stress does not target a single rate-limiting step but rather suppresses the entire glucosinolate biosynthesis cascade in a coordinated manner.

**Fig. 6.**
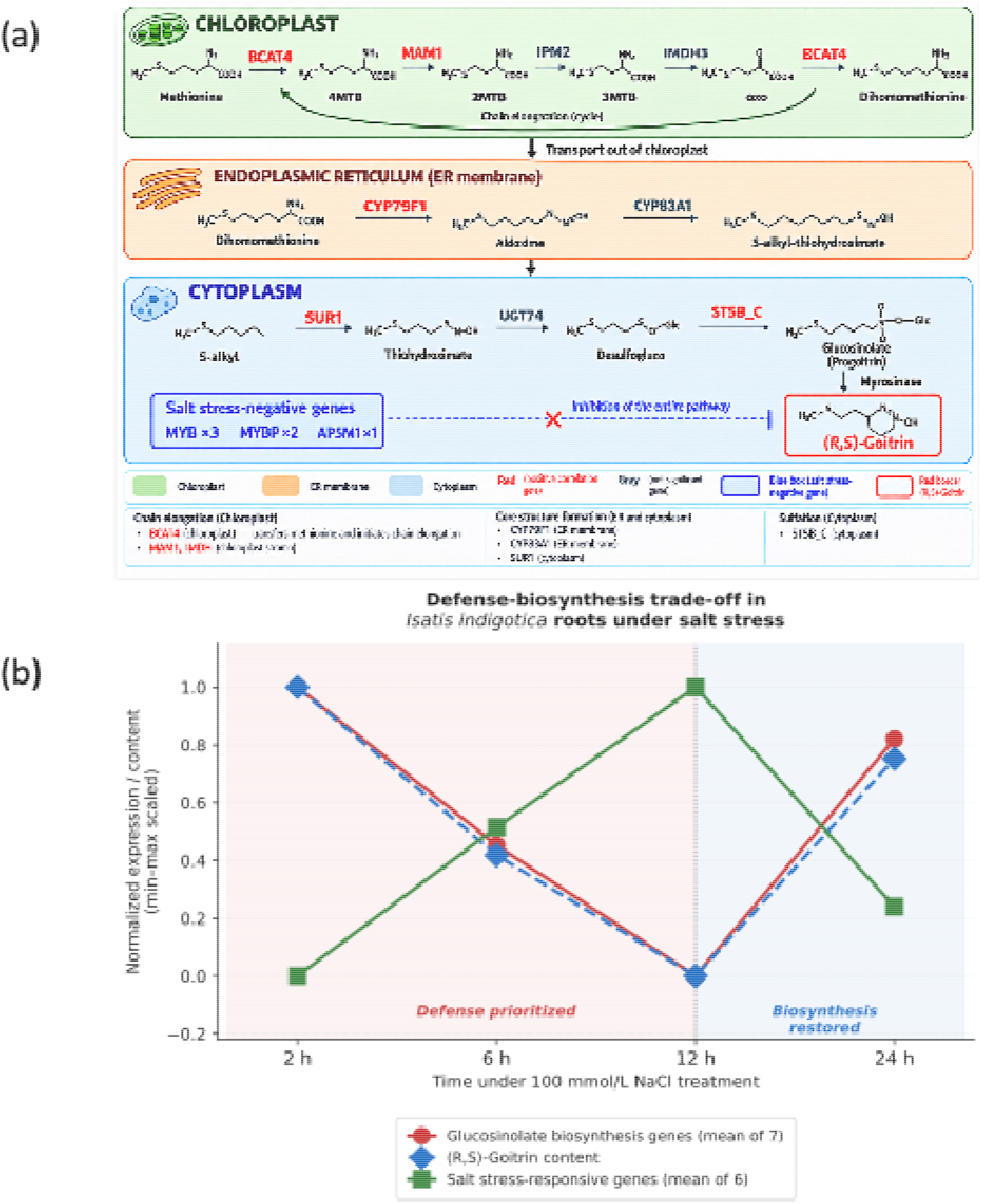
Proposed model of salt stress-mediated regulation of goitrin biosynthesis in *Isatis indigotica* roots. **(a)** Subcellular localization of the glucosinolate biosynthesis pathway across the chloroplast, endoplasmic reticulum, and cytoplasm. Gene names in red indicate significant positive correlation with goitrin accumulation (r > 0.957, p < 0.05); gray indicates other pathway genes. The blue dashed arrow (⊣) represents the proposed suppressive effect of salt stress-responsive genes (MYB ×3, MYBP ×2, AP5M1; blue box) on pathway activity, based on their negative correlation with goitrin content (p < 0.05). **(b)** Temporal dynamics of min–max normalized biosynthesis gene expression (mean TPM of 7 positively correlated genes, red), (R,S)-goitrin content (blue dashed), and stress-responsive gene expression (mean TPM of 6 negatively correlated genes, green) at 2, 6, 12, and 24 h under 100 mmol/L NaCl. The red-shaded region (2–12 h) indicates the defense prioritization phase; the blue-shaded region (12–24 h) indicates the partial recovery phase.

Concurrently, six salt stress-responsive genes, predominantly *MYB* and *MYBP* transcription factors together with *AP5M1*, showed expression dynamics inversely coupled to goitrin accumulation across the four time points. The opposing temporal trajectories of these two gene sets, with biosynthesis genes declining as stress-responsive genes rise during the early stress phase (2–12 h), and biosynthesis genes recovering as stress-responsive genes attenuate by 24 h, are illustrated in the proposed mechanistic model (**Fig. 6b**). This model depicts salt stress as triggering a transcriptional switch in which defense-related gene networks are prioritized at the expense of secondary metabolite biosynthesis, with the balance gradually shifting back toward biosynthesis as the plant begins to acclimate to the saline environment.

This transient suppression of goitrin biosynthesis is consistent with the growth–defense trade-off framework, which posits that plants under acute stress preferentially redirect limited metabolic resources toward immediate defense responses at the expense of secondary metabolite biosynthesis (He et al., 2022; Monson et al., 2022). As the stress response attenuates, secondary metabolic capacity is progressively restored, a pattern also observed in medicinal plants where the duration of stress exposure, rather than its intensity alone, determines the eventual trajectory of phytochemical accumulation (Taratima et al., 2025). It is noteworthy that glucosinolate levels have been reported to increase under salt stress in some *Brassica* species at comparable NaCl concentrations (Fatima et al., 2024), whereas the contrasting response observed in *I. indigotica* roots, a transient decline followed by partial recovery, suggests that the direction and magnitude of salt stress effects on glucosinolate accumulation may be species-specific, organ-specific, and highly dependent on the temporal scale of observation. The short-term time-series design employed in the present study revealed a dynamic suppression-recovery pattern that would be obscured in single time-point studies, underscoring the importance of temporal resolution in characterizing stress-responsive secondary metabolism in medicinal plants.

## 4. Conclusion

This study reveals that salt stress transiently suppresses (R,S)-goitrin accumulation in Isatis indigotica roots, with partial recovery as plants acclimate, contrary to the commonly reported pattern in which moderate abiotic stress promotes secondary metabolite accumulation. Time-series transcriptome analysis identified seven glucosinolate biosynthesis genes and six salt stress-responsive genes with opposing temporal expression dynamics, supporting a defense-biosynthesis trade-off model in which stress defense is prioritized over goitrin biosynthesis during the acute response phase. These findings provide new insight into the dynamic regulation of medicinal plant quality under salinity and identify transcriptional correlates of goitrin suppression that warrant further functional characterization.

## CRediT authorship contribution

Conceptualization: LH, YL, YZ^3^; Data curation: LH,YL; Formal analysis: LH, YZ^3^; Funding acquisition: HY; Investigation: YZ^1^; Methodology: LH, YL; Project administration: HY, YZ^1^; Resources: HY, YZ^1^; Supervision: HY, YZ^1^; Validation: LH, YL,YZ^3^; Visualization: LH,YL; Writing– original draft: LH; Writing– review & editing: LH, YL.

YZ^1^:Yueqiang Zhang. YZ^3^:Yanyan Zhang.

## Data availability statement

Data presented in this paper are available from the authors upon reasonable request.

